# The sodium/glucose cotransporter 2 inhibitor empagliflozin is a pharmacological chaperone of cardiac Na_v_1.5 channels

**DOI:** 10.1101/2025.05.15.654221

**Authors:** Jakob Sauer, Jessica Marksteiner, Martin Hohenegger, Hannes Todt, Helmut Kubista, Christopher Dostal, Attila Kiss, Bruno K. Podesser, Isabella Salzer, Xaver Koenig, Anna Stary-Weinzinger, Karlheinz Hilber, Oliver Kudlacek

## Abstract

Diminished peak sodium current (I_Na_) is a causative factor for slowed ventricular conduction and cardiac arrhythmias in patients with Duchenne muscular dystrophy (DMD), a devastating muscle disease triggered by dystrophin deficiency. Recently, we showed that chronic administration of the sodium/glucose cotransporter 2 (SGLT2) inhibitor empagliflozin (EMPA) restores diminished peak I_Na_ in ventricular cardiomyocytes from the dystrophin-deficient mdx mouse model of DMD. Here, we aimed to elucidate the underlying mechanism. Whole cell patch clamp studies revealed that 24 h incubation of dystrophic (mdx) ventricular cardiomyocytes with EMPA significantly increases peak I_Na_ in a concentration-dependent manner (EC_50_=94 nM). The enhancing effect on peak I_Na_ also occurred in dystrophic cardiac Purkinje fibers, Na_v_1.5-expressing tsA201 cells, as well as in dystrophic (DMD^mdx^) rat cardiomyocytes, and was also exerted by two other SGLT2 inhibitors. Immunofluorescence studies suggested that chronic EMPA treatment increases Na_v_1.5 plasma membrane expression. Peak I_Na_ enhancement by EMPA depended on functional anterograde trafficking of Na_v_1.5. The local anesthetic mexiletine, a well-known pharmacological chaperone of Na_v_1.5, enhanced peak I_Na_ in a similar manner as EMPA. Further, mutation of human Na_v_1.5 at a site important for local anesthetic binding (Y1767A) completely abolished the ability of both EMPA and mexiletine to enhance peak I_Na_. Finally, the importance of Y1767 for drug-induced modulation of peak I_Na_ was confirmed by molecular docking simulations. Our findings suggest that EMPA acts as a pharmacological chaperone of Na_v_1.5 channels. Its chronic administration may reduce arrhythmia vulnerability in patients with DMD and other arrhythmogenic pathologies associated with diminished peak I_Na_.

## 1. Introduction

In arrhythmia disorders associated with reduced peak sodium current (I_Na_), such as Brugada syndrome, the upstroke of the cardiac action potential and consequently electrical impulse conduction are slowed, thereby setting the stage for re-entrant arrhythmias and sudden cardiac death.^1^ A reduced cardiac peak I_Na_, impaired ventricular conduction, and concomitant arrhythmias are also features observed in patients with Duchenne muscular dystrophy (DMD)^2,3^ and animal models for this disease.^4–9^ No pharmacological treatments are currently available to enhance peak I_Na_ and restore cardiac conduction.

We recently showed^10^ that long-term treatment with therapeutic doses/concentrations of empagliflozin (EMPA), an inhibitor of sodium/glucose cotransporter 2 (SGLT2) in clinical use to treat type II diabetes and non-diabetic heart failure^11–13^, completely rescues abnormally reduced peak I_Na_ in ventricular cardiomyocytes from the most commonly used animal model for DMD, the dystrophin-deficient mdx mouse.^14^ The mechanism by which EMPA enhances peak I_Na_ in dystrophic myocytes, however, remained unknown.

Several groups recently reported that EMPA modulates the cardiac Na channel Na_v_1.5. In particular, application of the drug selectively inhibited the detrimental so-called “late I_Na_”, which is abnormally enhanced in ventricular cardiomyocytes derived from certain mouse models for heart failure.^15–17^ By simulations of EMPA docking to a three-dimensional homology model of human Na_v_1.5 and site-directed mutagenesis, Philippaert et al. showed that EMPA binds to Na_v_1.5 in the same region as local anesthetics.^15^ This interaction was crucial for inhibition of late I_Na_ by the drug. Local anesthetics such as lidocaine^18^ and mexiletine (MEXI)^19–21^ act as pharmacological chaperones of Na_v_ channels, thereby increasing their membrane trafficking. Disruption of the local anesthetic binding site by mutagenesis led to loss of Na_v_1.5 trafficking enhancement by MEXI^22^, suggesting that this drug has to bind to the Na_v_1.5 channel in order to be able to act as a pharmacochaperone.

Here, we used the mdx mouse as a model for an arrhythmia disorder associated with reduced peak I_Na_, and heterologous expression of wild-type and mutant human Na_v_1.5 channels to explore the following hypothesis: EMPA is a pharmacological chaperone of the cardiac sodium channel Na_v_1.5.

## 2. Methods

### 2.1 Ethical approval

The study conformed to the guiding principles of the Declaration of Helsinki and coincides with the rules of the Animal Welfare Committee of the Medical University of Vienna. The applied experimental protocols were approved by the Austrian Science Ministry. The respective ethics vote has the number BMWFW-66.009/0175-WF/V/3b/2015.

### 2.2 Animal models

Dystrophin-deficient mdx mice^14^ on the BL10 background (C57BL/10ScSn-Dmdmdx/J) in an age range between 16 and 26 weeks were used for isolation of ventricular cardiomyocytes of the working myocardium (termed ventricular cardiomyocytes throughout the text). This mouse line was originally purchased from Charles River Laboratories. For isolation of cardiac Purkinje fibers, 19-to 23-week-old double transgenic mdx-Cx40^eGFP/+^ mice were used. In these mice, enhanced green fluorescent protein (eGFP) is expressed under control of the connexin 40 (Cx40) gene.^8,23^ Dystrophin-deficient DMD^mdx^ Sprague Dawley rats^24^ originated from INSERM-CRTI UMR 1064 (Nantes). Cardiomyocytes were isolated when these rats had reached an age of 17 or 18 weeks. Only male animals were used in this study because of the X-linked inheritance of DMD and potential translational relevance to human patients as in Haffner et al.^25^ Genotyping of the animals was performed using standard PCR assays.

### 2.3 Drugs

Empagliflozin (EMPA), dapagliflozin (DAPA), sotagliflozin (SOTA), mexiletine (MEXI) hydrochloride and brefeldin A (BFA) were purchased from MedChemExpress. Cycloheximide (CHX) was purchased from Sigma-Aldrich. EMPA, DAPA, SOTA and BFA were dissolved in dimethyl sulfoxide (DMSO). MEXI hydrochloride and CHX were dissolved in H_2_O (Milli-Q^®^). Concentrations of the stock solutions were 100 mM (EMPA, DAPA, SOTA and MEXI hydrochloride) and 10 mg/ml (CHX and BFA). For experiments with cardiomyocytes, the final working concentrations were 1 µM (EMPA, DAPA and SOTA), 10 µM (MEXI hydrochloride), 5 µg/ml (BFA) and 50 µg/ml (CHX), unless otherwise noted. For experiments with tsA201 cells, the final working concentrations were 10 µM (EMPA) and 100 µM (MEXI hydrochloride). The use of a tenfold higher EMPA concentration in tsA201 cell compared to cardiomyocyte experiments was justified because of the presence of 10 % fetal bovine serum during the tsA201 cell incubation procedure and the extent of plasma protein binding of EMPA of ∼90 %.^26^ Thus, considering plasma protein binding, we estimated the free (effective) concentration of EMPA for incubation of tsA201 cells to amount to ∼1 µM. A similar calculation was performed in Dago et al.^27^

### 2.4 Isolation of cardiomyocytes and drug incubation procedure

Mdx mice, as well as DMD^mdx^ rats, were anesthetized with isoflurane (2 %, inhalation) and sacrificed by cervical dislocation. Ventricular cardiomyocytes of the working myocardium were then isolated from their hearts using a Langendorff setup (Hugo Sachs Elektronik, March, Germany), according to the procedure described in detail in our previous work.^5^ Cardiac Purkinje fibers were isolated from mdx-Cx40^eGFP/+^ mice using the same protocol. In brief, the heart was excised, and a cannula was inserted into the aorta for retrograde perfusion with Ca-free solution containing 0.17 mg/ml Liberase TH (Roche) at 37 °C for 10 min. Thereafter, the ventricles were pulled into pieces and incubated on a shaker at 37 °C. Subsequently, the Ca concentration was increased to 150 μM over 30 min in four steps. To liberate single cardiomyocytes, pieces of digested ventricular tissue were triturated. After a centrifugation step, the cells were resuspended in Minimum Essential Medium (MEM)-α, containing ITS media supplement (diluted 1:100), 2 mM L-glutamine, 100 U/ml penicillin, 0.1 mg/ml streptomycin and 17 μM blebbistatin (Sigma-Aldrich). For patch clamp experiments, cardiomyocytes were then plated on Matrigel (Corning)-coated 3.5 cm culture dishes. For immunofluorescence stainings, cardiomyocytes were plated on Matrigel-coated glass cover slips. For drug incubation experiments, isolated cardiomyocytes in cell culture medium were exposed to various drugs, or combinations thereof, for a duration of 24 h. The only exception here was cardiomyocyte incubation with CHX. This drug is strongly cell toxic, and, therefore, incubation time was limited to 4 h. Prior to any performed experiment, drugs were removed from the bathing solutions. Untreated control cardiomyocytes were exposed to respective concentrations of the solvent for the same duration as treated cardiomyocytes to the drug. DMSO concentrations were equal in all compared experimental groups. Drug-treated cardiomyocytes and untreated control cells always originated from the same cardiomyocyte isolation (identical animal). During the incubation period, the cardiomyocytes were maintained at a temperature of 37 °C in a humidified atmosphere of 5 % CO_2_.

### 2.5 Mutagenesis

For heterologous expression of human Na_v_1.5 fused to GFP at its C-terminus, a plasmid first described by Zimmer et al.^28^ was used. Point mutations for Na_v_1.5 (F1760A and Y1767A) were introduced using the QuikChange Lightning site-directed mutagenesis kit (Agilent, Vienna, Austria), according to the manufacturer’s protocol. Primers for mutagenesis were designed using the online tool from Agilent (agilent.com) and synthesized by Microsynth AG (Balgach, Switzerland). Mutagenesis was confirmed by sequencing the entire coding region at LGC Genomics GmbH (Berlin, Germany). The following primers were used for mutagenesis:

Na_v_1.5-F1760A fw: CACCTACATCATCATCTCCGCCCTCATCGTGGTCAACATG

Na_v_1.5-F1760A rv: CATGTTGACCACGATGAGGGCGGAGATGATGATGTAGGTG

Na_v_1.5-Y1767A fw: CCTCATCGTGGTCAACATGGCCATTGCCATCATCCTGGAG

Na_v_1.5-Y1767A rv: CTCCAGGATGATGGCAATGGCCATGTTGACCACGATGAGG

### 2.6 Culture, transient transfection and drug incubation of tsA201 cells

tsA201 cells (ECACC no. 96121229; Salisbury, United Kingdom; RRID: CVCL_2737) were cultured in Dulbecco’s Modified Eagle Medium (high glucose) supplemented with 10 % fetal bovine serum (Capricorn Scientific) at a temperature of 37 °C in a humidified atmosphere of 5 % CO_2_. For transfection, they were seeded on 3.5 cm dishes. 48 h before the patch clamp experiments, the cells were transiently transfected with human Na_v_1.5 wild-type (wt), F1760A or Y1767A using a custom-made protocol. Per 3.5 cm dish, 1 µg plasmid DNA was diluted in 55 µl of a 150 mM NaCl solution. Thereafter, 20 μl of 10 μM polyethylenimine (Polysciences, Inc., Hirschberg an der Bergstrasse, Germany) were added, and the mixture was vortexed for 10 s. After allowing it to rest for 15 min at room temperature, the mixture was applied to the cells. 24 h after transfection, the cells were split. Half of the dishes obtained by cell splitting were exposed to the drug (EMPA or MEXI) for 24 h. Prior to any performed experiment, drugs were removed from the bathing solutions. Cells of the control group, which originated from the same transfected dish as the drug-treated cells, were incubated with a respective concentration of the solvent for the same duration. During the incubation period, the cells were maintained at 37 °C and 5 % CO_2_.

### 2.7 Sodium current recordings

The whole cell patch clamp technique was used to record I_Na_ in isolated cardiomyocytes and transiently transfected tsA201 cells. All recordings were performed at room temperature (22 ± 1.5 °C) using an Axopatch 200B patch clamp amplifier, a Digidata1440 digitizer and the software Clampex 10.7 (Axon Instruments, Union City, CA, USA). Measurements were performed after the drug incubation period. Before the start of the electrophysiological experiments, cells were washed with bath solution in order to remove the drugs used for incubation. Ventricular cardiomyocytes and Purkinje fibers were bathed in a solution that consisted of (in mM) 5 NaCl, 135 N-methyl-D-glucamine, 1 CaCl_2_, 1 MgCl_2_ and 10 HEPES, adjusted to pH 7.4 with HCl. This bath solution additionally contained 17 µM blebbistatin. tsA201 cells were bathed in a solution consisting of (in mM) 15 NaCl, 125 N-methyl-D-glucamine, 1 CaCl_2_, 1 MgCl_2_ and 10 HEPES, adjusted to pH 7.4 with HCl. During the recordings, patched cells were continuously superfused with fresh bath solution via a DAD-8-VC superfusion system (ALA Scientific Instruments, Westbury, NY, USA). Patch pipettes were formed from aluminosilicate glass capillaries (A120-77-10; Science Products, Hofheim, Germany) with a P-97 horizontal puller (Sutter Instruments, Novato, CA, USA). When filled with pipette solution, they had resistances between 1 and 1.4 MΩ (for recordings in cardiomyocytes) or between 1.8 and 2.2 MΩ (for recordings in tsA201 cells). The pipette solution contained (in mM) 5 NaCl, 110 CsF, 10 EGTA and 10 HEPES, adjusted to pH 7.3 with CsOH. Only single cardiomyocytes or single fluorescent tsA201 cells were patched. Purkinje fibers were discriminated from ventricular cardiomyocytes by their typical morphology and eGFP signal, as previously described.^8^ A holding potential of −116 mV (cardiomyocytes) or −146 mV (tsA201 cells) was chosen to guarantee full Na channel availability. To measure current density-voltage relationships, I_Na_ was activated by 25 ms depolarizing voltage steps ranging from −96 to −6 mV (cardiomyocytes) or from −116 to 14 mV (tsA201 cells). All membrane voltages were corrected for liquid junction potentials. Leak currents and capacitive transients were subtracted using a p/4 protocol. Data were low-pass filtered with 10 kHz and digitized at 50 kHz. Data analysis was performed with Clampfit 10.7 (Axon Instruments) and GraphPad Prism 8 (San Diego, CA, USA) software. Peak I_Na_ amplitudes at the various voltage steps were determined. These values were divided by the membrane capacitance to yield current densities, which were then plotted against the test pulse voltages. Data were then fit with the function: I=G_max_·(V-V_rev_)/(1+exp((V_50_-V)/K)), where I is the current, G_max_ is the maximal conductance, V is the membrane potential, V_rev_ is the reversal potential, V_50_ is the voltage at which the half-maximal activation occurred, and K is the slope factor. For construction of a concentration-response curve, current densities at −41 mV were plotted against the EMPA concentration. Data were then fit with the function: Y=Bottom+(Top-Bottom)/(1+10^((LogEC_50_-X)·HillSlope)), where EC_50_ is the concentration at which the half-maximal effect was reached.

### 2.8 Immunofluorescence

Immunofluorescence stainings of isolated mdx ventricular cardiomyocytes were performed after the 24 h incubation period with EMPA or solvent. First, the culture medium was removed from the glass cover slips and the cells were washed with PBS. Then, they were fixed with 4 % paraformaldehyde for 10 min at room temperature. After three washing steps with PBS, the cardiomyocytes were permeabilized with 0.05 % Triton X-100 for 5 min at room temperature. Thereafter, they were washed three times with PBS and then incubated with 10 % horse serum for 1 h at room temperature, followed by three more washing steps with PBS. Subsequently, the cells were incubated with the ASC-005 anti-Na_v_1.5 rabbit primary antibody (Alomone Labs, diluted 1:100 in PBS) for 1 h at room temperature. This antibody was chosen because it had previously been successfully used by others in immunofluorescence and western blot studies with cardiomyocytes.^4,20,29^ After incubation with the primary antibody, the cells were washed three times with PBS and then exposed to a donkey anti-rabbit secondary antibody conjugated to Alexa Fluor Plus 555 (# A32794, Thermo Fisher Scientific; diluted 1:500 in PBS) for 1 h at room temperature. Finally, after three more washing steps with PBS, the glass cover slips harboring the cardiomyocytes were mounted on microscope slides. The slides were then dried and stored at 4 °C. Immunofluorescence images were acquired using a Nikon A1R confocal laser scanning microscope with a 60x oil objective and NIS-Elements AR software. Alexa Fluor Plus 555 was excited at 561 nm and emitted light was collected at 595 nm. Microscope settings were equal for all images. Using these settings, no auto- or Alexa Fluor Plus 555 fluorescence was visible for negative control cardiomyocytes, which had been incubated only with the secondary antibody (no primary antibody). For quantification with NIS-Elements AR, one region of interest was manually drawn for each individual cardiomyocyte. This region of interest always included parts of the lateral membrane and intercalated disc region and was drawn in the transmitted light image (see Figure 2C), thereby ensuring an unbiased selection of the areas. Then, the mean Alexa Fluor Plus 555 fluorescence intensity within the region of interest was measured and used as an indicator for Na_v_1.5 plasma membrane expression.

### 2.9 Molecular modeling

The 3D conformers of empagliflozin (EMPA), mexiletine (MEXI), dapagliflozin (DAPA) and sotagliflozin (SOTA) were prepared with Ligandscout 4.4.8 (Inte:Ligand GmbH, Vienna, Austria). This included enantiomer and tautomer generation, energy minimization with the MMFF94 force field and the generation of 25 conformers per drug. The cryo-EM structure of rat and human Na_v_1.5 channels (PDB ID: **7XSU**, resolution 3.4Å, **8T6L**, 3.3Å, PDB ID: **6UZ0**, 3.24Å, PDB ID: **6UZ3**, 3.5Å, PDB ID: **7FBS**, 3.4Å, PDB ID: **7K18**, 3.3Å, PDB ID: **8F6P**, 3.2Å, PDB ID: **6LQA**, 3.3Å and PDB ID: **7DTC**, 3.3Å) were obtained from the RCSB Protein Data Bank, and structure quality was assessed using the **wwPDB Validation** 3D report, available via RCSB (https://www.rcsb.org). In addition, a model of Na_v_1.5 was generated using the Alphafold3 server^30^, with UniProt entry Q14524 SCN5A_HUMAN. Docking was performed using GOLD version 2020.2.^31^ Each docking underwent 125,000 genetic algorithm operations. Y1767 and F1760, as well as all residues within a radius of 10 Å were selected as binding pocket. The scoring function used was CHEMPLP^32^, and the top 10 highest-ranking poses out of 100 were visually inspected. Consensus docking figures were prepared with PyMol v2.5.5 (Pymol, Schrodinger LCC).

### 2.10 Statistical data analysis

Comparisons were made using a nested analysis respecting the hierarchical data structure (measurements of n cells from m animals or transfections) described by Sikkel et al.^33^ Since drug-treated and untreated control cells always originated from the same animal or transfection, hierarchical testing was paired. A P value < 0.05 was considered to indicate statistical significance. Data points represent means ± SE.

## 3. Results

### 3.1 EMPA enhances peak I_Na_ in dystrophin-deficient cardiomyocytes and Na_v_1.5-expressing tsA201 cells

We recently reported that peak I_Na_ of ventricular cardiomyocytes derived from mdx mice, which had received clinically relevant doses of EMPA for 4 weeks, was fully restored to wild-type level.^10^ Furthermore, 24 h incubation of isolated mdx cardiomyocytes with 1 µM EMPA significantly increased their peak I_Na_. These findings implied that chronic EMPA treatment completely rescues abnormally reduced peak I_Na_ of dystrophin-deficient ventricular cardiomyocytes. EMPA treatment *in vivo* and *in vitro* did not affect the Na channel gating properties, and acute application of EMPA had no impact on peak I_Na_ of mdx cardiomyocytes.^10^

Figure 1A shows original I_Na_ traces, elicited by the pulse protocol displayed in the inset, which were recorded from ventricular cardiomyocytes isolated from dystrophin-deficient mdx mice. The myocytes had either been untreated (control, DMSO), or incubated for 24 h with 0.1 µM or 1 µM EMPA. Peak current density-voltage relationships, derived from a series of such experiments (Fig. 1B), showed that EMPA increases peak I_Na_ densities in a concentration-dependent manner. The respective Na channel activation parameters were unaltered (Table 1). A concentration-response curve is shown in Figure 1C, and fitting of the data (for function see Methods) revealed an EC_50_ value of 94 nM (Hill Coeff., 1.3). Together, these data suggested that chronic EMPA treatment increases peak I_Na_ of dystrophic ventricular cardiomyocytes, and this effect is concentration-dependent. At 1 µM EMPA, a therapeutically relevant concentration^15,16,27^, the effect had reached its maximum (Fig. 1C).

**Figure 1.**
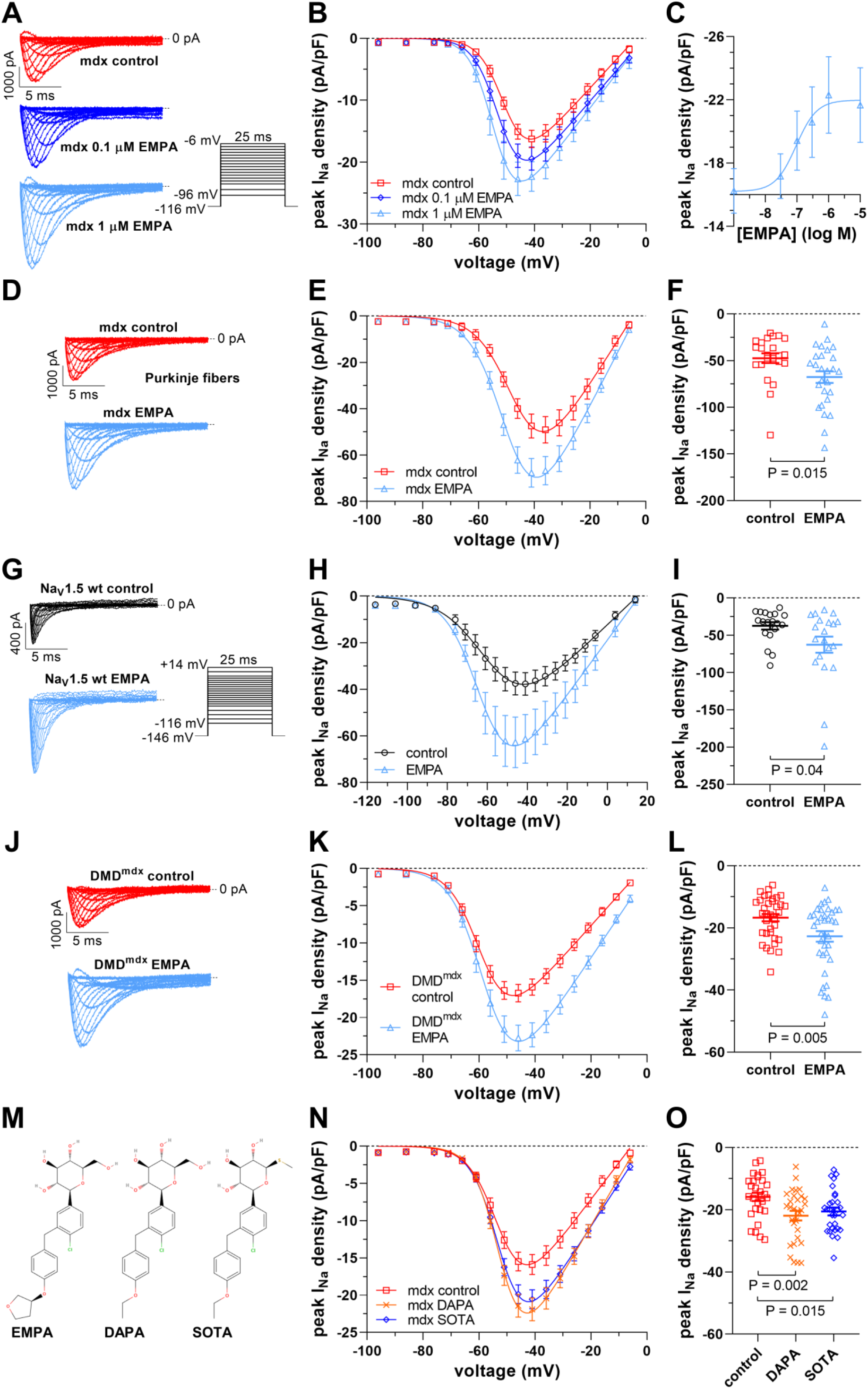
EMPA and other SGLT2 inhibitors enhance peak I_Na_ densities in dystrophin-deficient cardiomyocytes and Na_v_1.5-expressing tsA201 cells. **A-C**, Concentration-response relationship of peak I_Na_ enhancement by 24 h EMPA incubation in ventricular cardiomyocytes derived from mdx mice. **A**, Typical original I_Na_ traces of an mdx cardiomyocyte after 24 h incubation under control conditions and after 24 h incubation with 0.1 µM or 1 µM EMPA, elicited by the pulse protocol displayed in the inset. **B**, From a series of such experiments [n = 12 cells (mdx control), 12 cells (mdx 0.1 µM EMPA), and 13 cells (mdx 1 µM EMPA); all cells originating from the same 3 mdx hearts], peak I_Na_ density-voltage relationships were derived. Data points are represented as means ± SE. Parameters for I_Na_ activation derived from fits of the current-voltage relationships (function described in Methods) are given in Table 1. **C**, Peak I_Na_ densities at −41 mV were plotted against the EMPA concentration. Data were fit with a function given in the Methods and yielded an EC_50_ value of 94 nM (Hill Coeff., 1.3). **D-F**, Effect of 24 h incubation with 1 µM EMPA on peak I_Na_ of cardiac Purkinje fibers derived from mdx-Cx40^eGFP/+^ mice. **D**, Typical original current traces of an mdx Purkinje fiber after incubation under control conditions or after EMPA treatment. **E**, Respective peak I_Na_ density-voltage relationships [n = 23 cells (mdx control) and 27 cells (mdx EMPA); all cells originating from the same 4 mdx hearts]. **F**, Dot plot comparing the maximum peak I_Na_ densities of untreated control and EMPA-treated mdx Purkinje fibers at −41 mV. A significant difference existed between control and EMPA-treated cells. **G-I**, Effect of 24 h incubation with 10 µM EMPA (estimated free EMPA concentration: 1 µM) on peak I_Na_ of tsA201 cells expressing wild-type Na_v_1.5. **G**, Typical original current traces of a Na_v_1.5-expressing tsA201 cell after incubation under control conditions or after EMPA treatment (for pulse protocol see inset). **H**, Respective peak I_Na_ density-voltage relationships [n = 19 cells (control) and 20 cells (EMPA); all cells originating from the same 4 transfections]. **I**, Dot plot comparing the maximum peak I_Na_ densities of untreated control and EMPA-treated Na_v_1.5-expressing tsA201 cells at −46 mV. A significant difference existed between control and EMPA-treated cells. **J-L**, Effect of 24 h incubation with 1 µM EMPA on peak I_Na_ of ventricular cardiomyocytes derived from DMD^mdx^ rats. **J**, Typical original current traces of a DMD^mdx^ rat cardiomyocyte after incubation under control conditions or after EMPA treatment. **K**, Respective peak I_Na_ density-voltage relationships [n = 35 cells (DMD^mdx^ control) and 37 cells (DMD^mdx^ EMPA); all cells originating from the same 4 DMD^mdx^ hearts]. **L**, Dot plot comparing the maximum peak I_Na_ densities of untreated control and EMPA-treated DMD^mdx^ cardiomyocytes at −46 mV. A significant difference existed between control and EMPA-treated cells. **M-O**, Effect of 24 h incubation with other SGLT2 inhibitors on peak I_Na_ of ventricular cardiomyocytes derived from mdx mice. **M**, Chemical structure comparison of EMPA (PubChem CID 11949646), dapagliflozin (DAPA, PubChem CID 9887712), and sotagliflozin (SOTA, PubChem CID 24831714). **N**, Peak I_Na_ density-voltage relationships from untreated control mdx cardiomyocytes [n = 28 cells], mdx myocytes treated with 1 µM DAPA [n = 30 cells], and mdx myocytes treated with 1 µM SOTA [n = 29 cells, all cells originating from the same 4 mdx hearts]. **O**, Dot plot comparing the maximum peak I_Na_ densities of untreated control, DAPA-treated, and SOTA-treated mdx cardiomyocytes at −41 mV. A significant difference existed between control and DAPA-treated cells, as well as between control and SOTA-treated cells.

**Table 1.**
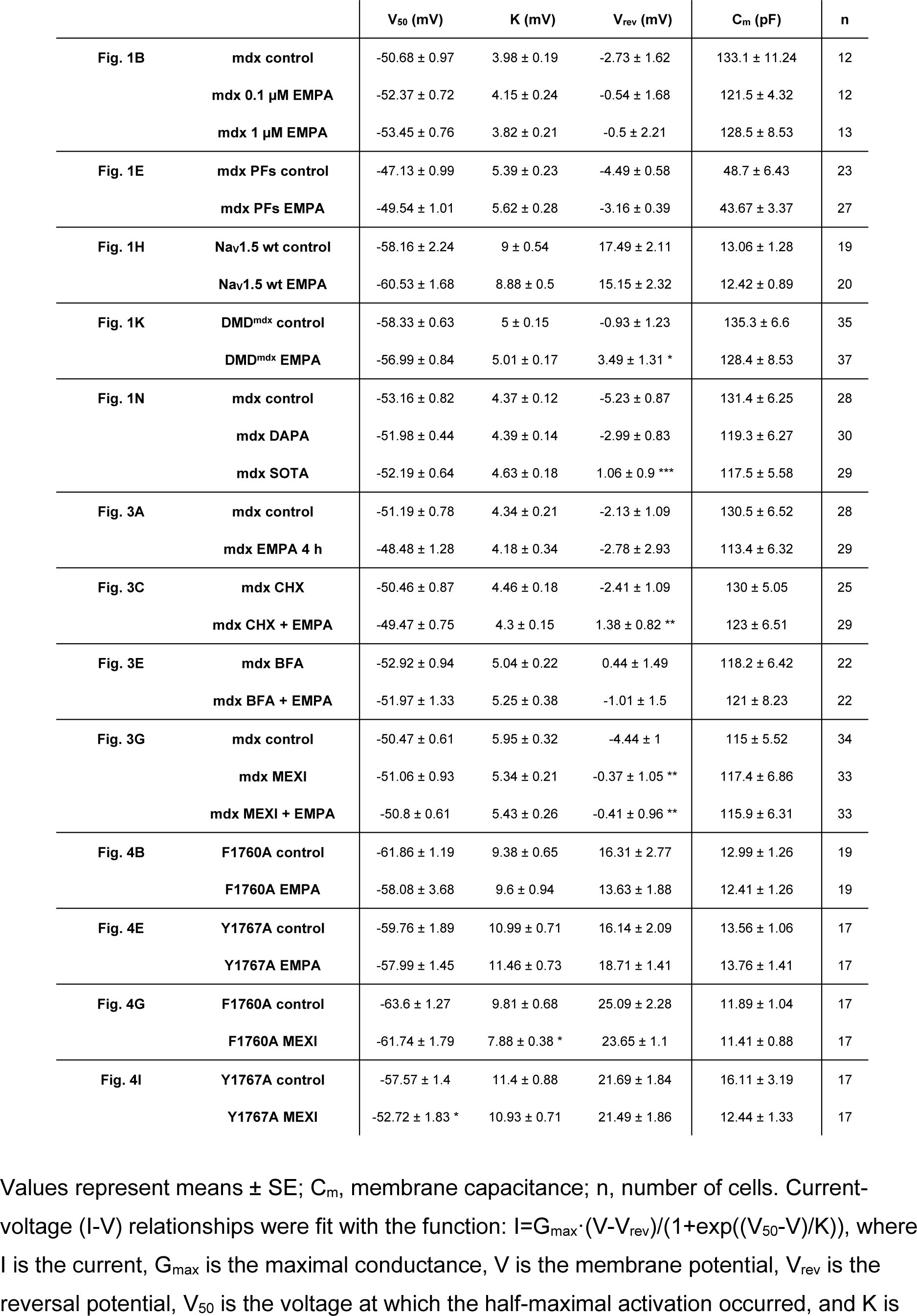
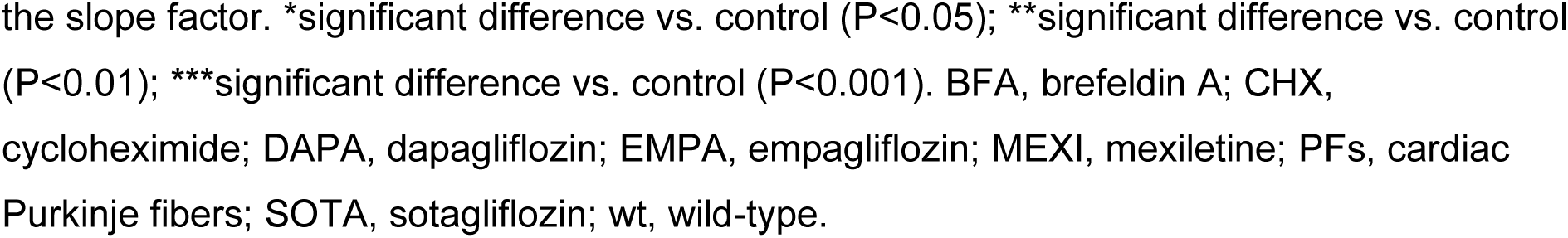
Parameters of I_Na_ activation in mdx cardiomyocytes and Na_v_1.5-expressing tsA201 cells.

| | | $V_{50}$ (mV) | K (mV) | $V_{rev}$ (mV) | $C_m$ (pF) | n |
| --- | --- | --- | --- | --- | --- | --- |
| Fig. 1B | mdx control | $-50.68 \pm 0.97$ | $3.98 \pm 0.19$ | $-2.73 \pm 1.62$ | $133.1 \pm 11.24$ | 12 |
| | mdx 0.1 μM EMPA | $-52.37 \pm 0.72$ | $4.15 \pm 0.24$ | $-0.54 \pm 1.68$ | $121.5 \pm 4.32$ | 12 |
| | mdx 1 μM EMPA | $-53.45 \pm 0.76$ | $3.82 \pm 0.21$ | $-0.5 \pm 2.21$ | $128.5 \pm 8.53$ | 13 |
| Fig. 1E | mdx PFs control | $-47.13 \pm 0.99$ | $5.39 \pm 0.23$ | $-4.49 \pm 0.58$ | $48.7 \pm 6.43$ | 23 |
| | mdx PFs EMPA | $-49.54 \pm 1.01$ | $5.62 \pm 0.28$ | $-3.16 \pm 0.39$ | $43.67 \pm 3.37$ | 27 |
| Fig. 1H | $Na_v1.5$ wt control | $-58.16 \pm 2.24$ | $9 \pm 0.54$ | $17.49 \pm 2.11$ | $13.06 \pm 1.28$ | 19 |
| | $Na_v1.5$ wt EMPA | $-60.53 \pm 1.68$ | $8.88 \pm 0.5$ | $15.15 \pm 2.32$ | $12.42 \pm 0.89$ | 20 |
| Fig. 1K | DMD<sup>mdx</sup> control | $-58.33 \pm 0.63$ | $5 \pm 0.15$ | $-0.93 \pm 1.23$ | $135.3 \pm 6.6$ | 35 |
| | DMD<sup>mdx</sup> EMPA | $-56.99 \pm 0.84$ | $5.01 \pm 0.17$ | $3.49 \pm 1.31^*$ | $128.4 \pm 8.53$ | 37 |
| Fig. 1N | mdx control | $-53.16 \pm 0.82$ | $4.37 \pm 0.12$ | $-5.23 \pm 0.87$ | $131.4 \pm 6.25$ | 28 |
| | mdx DAPA | $-51.98 \pm 0.44$ | $4.39 \pm 0.14$ | $-2.99 \pm 0.83$ | $119.3 \pm 6.27$ | 30 |
| | mdx SOTA | $-52.19 \pm 0.64$ | $4.63 \pm 0.18$ | $1.06 \pm 0.9^{***}$ | $117.5 \pm 5.58$ | 29 |
| Fig. 3A | mdx control | $-51.19 \pm 0.78$ | $4.34 \pm 0.21$ | $-2.13 \pm 1.09$ | $130.5 \pm 6.52$ | 28 |
| | mdx EMPA 4 h | $-48.48 \pm 1.28$ | $4.18 \pm 0.34$ | $-2.78 \pm 2.93$ | $113.4 \pm 6.32$ | 29 |
| Fig. 3C | mdx CHX | $-50.46 \pm 0.87$ | $4.46 \pm 0.18$ | $-2.41 \pm 1.09$ | $130 \pm 5.05$ | 25 |
| | mdx CHX + EMPA | $-49.47 \pm 0.75$ | $4.3 \pm 0.15$ | $1.38 \pm 0.82^{**}$ | $123 \pm 6.51$ | 29 |
| Fig. 3E | mdx BFA | $-52.92 \pm 0.94$ | $5.04 \pm 0.22$ | $0.44 \pm 1.49$ | $118.2 \pm 6.42$ | 22 |
| | mdx BFA + EMPA | $-51.97 \pm 1.33$ | $5.25 \pm 0.38$ | $-1.01 \pm 1.5$ | $121 \pm 8.23$ | 22 |
| Fig. 3G | mdx control | $-50.47 \pm 0.61$ | $5.95 \pm 0.32$ | $-4.44 \pm 1$ | $115 \pm 5.52$ | 34 |
| | mdx MEXI | $-51.06 \pm 0.93$ | $5.34 \pm 0.21$ | $-0.37 \pm 1.05^{**}$ | $117.4 \pm 6.86$ | 33 |
| | mdx MEXI + EMPA | $-50.8 \pm 0.61$ | $5.43 \pm 0.26$ | $-0.41 \pm 0.96^{**}$ | $115.9 \pm 6.31$ | 33 |
| Fig. 4B | F1760A control | $-61.86 \pm 1.19$ | $9.38 \pm 0.65$ | $16.31 \pm 2.77$ | $12.99 \pm 1.26$ | 19 |
| | F1760A EMPA | $-58.08 \pm 3.68$ | $9.6 \pm 0.94$ | $13.63 \pm 1.88$ | $12.41 \pm 1.26$ | 19 |
| Fig. 4E | Y1767A control | $-59.76 \pm 1.89$ | $10.99 \pm 0.71$ | $16.14 \pm 2.09$ | $13.56 \pm 1.06$ | 17 |
| | Y1767A EMPA | $-57.99 \pm 1.45$ | $11.46 \pm 0.73$ | $18.71 \pm 1.41$ | $13.76 \pm 1.41$ | 17 |
| Fig. 4G | F1760A control | $-63.6 \pm 1.27$ | $9.81 \pm 0.68$ | $25.09 \pm 2.28$ | $11.89 \pm 1.04$ | 17 |
| | F1760A MEXI | $-61.74 \pm 1.79$ | $7.88 \pm 0.38^*$ | $23.65 \pm 1.1$ | $11.41 \pm 0.88$ | 17 |
| Fig. 4I | Y1767A control | $-57.57 \pm 1.4$ | $11.4 \pm 0.88$ | $21.69 \pm 1.84$ | $16.11 \pm 3.19$ | 17 |
| | Y1767A MEXI | $-52.72 \pm 1.83^*$ | $10.93 \pm 0.71$ | $21.49 \pm 1.86$ | $12.44 \pm 1.33$ | 17 |

Values represent means ± SE; $C_m$ , membrane capacitance; n, number of cells. Current-voltage (I-V) relationships were fit with the function: $I = G_{max} \cdot (V - V_{rev}) / (1 + \exp((V_{50} - V)/K))$ , where I is the current, $G_{max}$ is the maximal conductance, V is the membrane potential, $V_{rev}$ is the reversal potential, $V_{50}$ is the voltage at which the half-maximal activation occurred, and K is
the slope factor. \*significant difference vs. control ( $P<0.05$ ); \*\*significant difference vs. control ( $P<0.01$ ); \*\*\*significant difference vs. control ( $P<0.001$ ). BFA, brefeldin A; CHX, cycloheximide; DAPA, dapagliflozin; EMPA, empagliflozin; MEXI, mexiletine; PFs, cardiac Purkinje fibers; SOTA, sotagliflozin; wt, wild-type.

We next tested if 24 h incubation with EMPA increased peak I_Na_ in cardiac Purkinje fibers, myocytes specialized for rapid ventricular impulse conduction, in a similar manner as in ventricular cardiomyocytes of the working myocardium. Figure 1D-F shows that EMPA (1 µM) enhanced peak I_Na_ densities in mdx Purkinje fibers by 43 % (Fig. 1F). I_Na_ activation parameters were again unaltered (Table 1). Comparison with our earlier work^8,9^ suggested that EMPA treatment restores I_Na_ in dystrophic Purkinje fibers to wild-type level.

Next, it was of interest if 24 h EMPA incubation also affected peak I_Na_ in tsA201 cells heterologously expressing human wild-type Na_v_1.5 channels. Figure 1G-I shows that EMPA (10 µM, estimated free concentration: 1 µM; see Methods) enhanced peak I_Na_ densities in Na_v_1.5-expressing tsA201 cells (for Na_v_1.5 channel activation parameters see Table 1). In spite of a large variation within the data due to heterologous expression of Na_v_1.5, a significant difference was found at −46 mV, the potential at which peak I_Na_ was maximum (Fig. 1I).

In order to exclude that the effect of EMPA on peak I_Na_ in dystrophic ventricular cardiomyocytes was a mouse-specific phenomenon, here, besides mdx mice, we also used a rat model of DMD, the DMD^mdx^ rat.^24,34^ Figure 1J-L clearly shows that, similar as in mouse mdx cardiomyocytes, 24 h EMPA incubation significantly enhanced the peak I_Na_ density in DMD^mdx^ rat myocytes. This finding, together with Dago et al., who reported that 24 h EMPA incubation increased peak I_Na_ in cardiomyocytes derived from human induced pluripotent stem cells^27^, excluded that EMPA’s effect on peak I_Na_ is only species-specific.

### 3.2 Dapagliflozin and sotagliflozin enhance peak I_Na_ in dystrophin-deficient ventricular cardiomyocytes

Next, we tested if the effect of EMPA on peak I_Na_ in dystrophic ventricular cardiomyocytes can also be generated by other SGLT2 inhibitors. Therefore, we used dapagliflozin (DAPA) and sotagliflozin (SOTA, a dual inhibitor of both SGLT1 and SGLT2^35^) in 24 h incubation experiments with mdx cardiomyocytes. Figure 1M-O shows that both DAPA (1 µM) and SOTA (1 µM) significantly enhanced peak I_Na_ densities in a similar manner to EMPA. This suggested that peak I_Na_ enhancement in dystrophic cardiomyocytes triggered by chronic EMPA treatment is not an effect specific for this particular drug, but is generated by SGLT2 inhibitors in general.

### 3.3 EMPA increases peak I_Na_ by enhancement of Na_v_1.5 channel trafficking to the plasma membrane

EMPA-induced peak I_Na_ enhancement in dystrophic cardiomyocytes may be caused by an increased number of functional Na channels in the plasma membrane. To test this hypothesis, we performed immunofluorescence experiments. These studies revealed that treatment of mdx ventricular cardiomyocytes with 1 µM EMPA for 24 h significantly enhances the mean fluorescence intensity along the plasma membrane (Fig. 2A-D). This suggested enhanced plasma membrane expression of Na_v_1.5 channels, which accords with an increase in the peak I_Na_ density after respective EMPA treatment.

**Figure 2.**
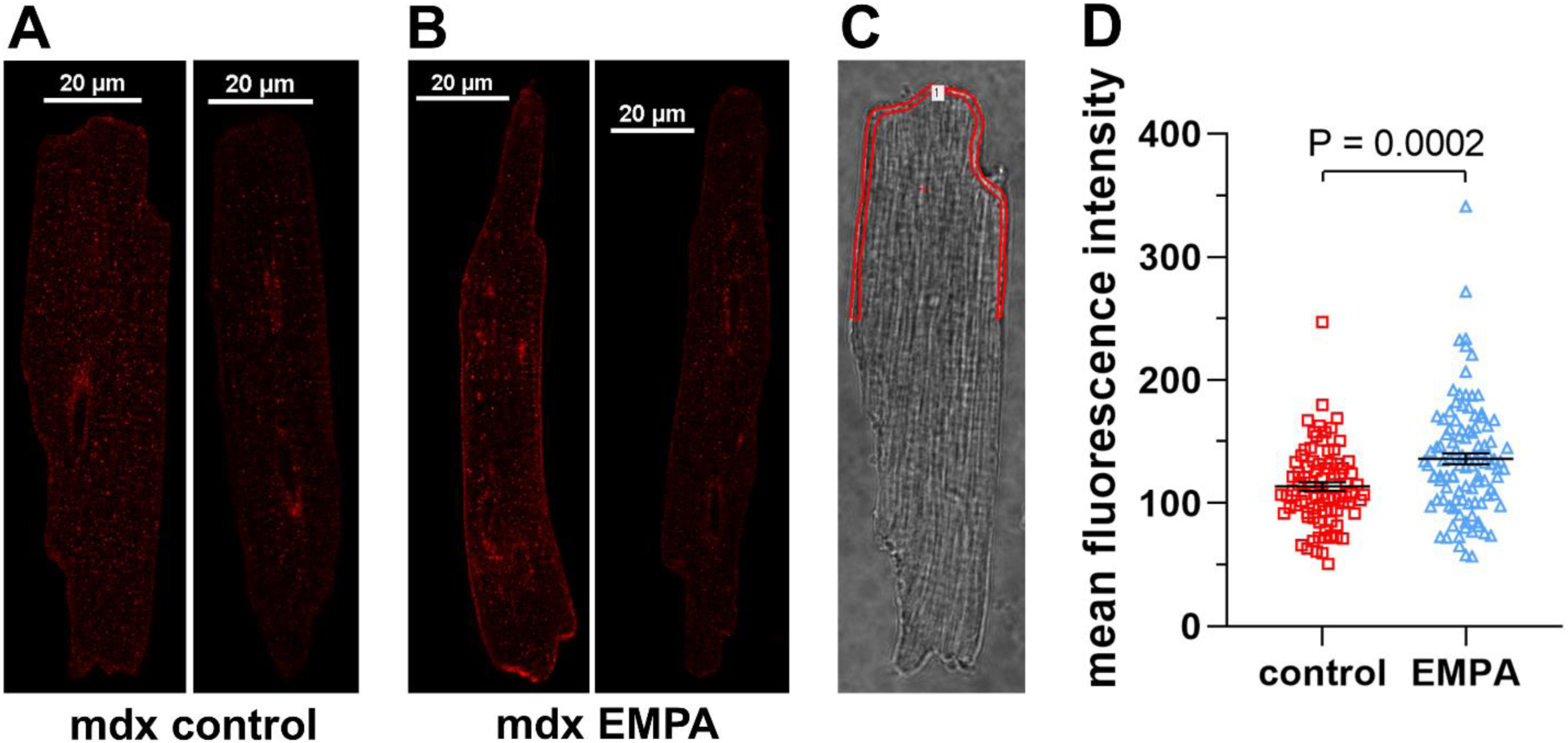
Chronic exposure to EMPA increases Na_v_1.5 plasma membrane expression in cardiomyocytes from dystrophin-deficient mdx mice. **A**, Representative confocal immunofluorescence images of isolated mdx ventricular cardiomyocytes stained using a Na_v_1.5-specific rabbit primary antibody (ASC-005, Alomone Labs; dilution 1:100) and a donkey anti-rabbit secondary antibody conjugated to Alexa Fluor Plus 555 (# A32794, Thermo Fisher Scientific; dilution 1:500). Staining of these two cells was performed after a 24 h incubation under control conditions. **B**, Representative images of two mdx cardiomyocytes stained after a 24 h incubation with 1 µM EMPA. **C**, Exemplary region of interest for quantification of fluorescence, always drawn in the transmitted light image in order to guarantee an unbiased selection of the areas. The mean fluorescence intensity (in arbitrary units) within the region of interest (one for each cardiomyocyte) was used as a marker for Na_v_1.5 expression. **D**, Mean fluorescence intensities for control and EMPA-treated cardiomyocytes from 3 mdx hearts [n = 82 cells (mdx control) and 104 cells (mdx EMPA)]. A significant difference existed between control and EMPA-treated cells. Each dot represents one cell. In addition, means ± SE are shown.

EMPA-induced enhancement of Na_v_1.5 plasma membrane expression could be generated by promotion of anterograde Na_v_1.5 channel trafficking by the drug. Because EMPA can directly bind to Na_v_1.5^15^, this could occur via pharmacological chaperoning. Thus, EMPA binding may facilitate Na_v_1.5 protein folding and export from the endoplasmic reticulum (ER), thereby allowing more channels to be trafficked to the plasma membrane. To test this hypothesis, we performed a series of experiments described in the following.

First, we used the protein synthesis inhibitor cycloheximide (CHX) and the anterograde trafficking inhibitor brefeldin A (BFA) to test under which conditions EMPA can exert its effect on peak I_Na_ in dystrophic cardiomyocytes. Figure 3A-D shows that the presence of CHX during a 4 h cardiomyocyte incubation period did not have an obvious effect on peak I_Na_ enhancement by EMPA. The presence of BFA during a 24 h incubation period, on the other hand, completely impeded the effect of EMPA (Fig. 3E-F). Together, these findings suggested that *de novo* protein synthesis of Na_v_1.5 (inhibited by CHX) is not required for EMPA-induced enhancement of peak I_Na_ in dystrophic cardiomyocytes, but in fact the transport of newly synthesized Na channels from the ER to the Golgi apparatus (inhibited by BFA) is essential for EMPA to be effective.

**Figure 3.**
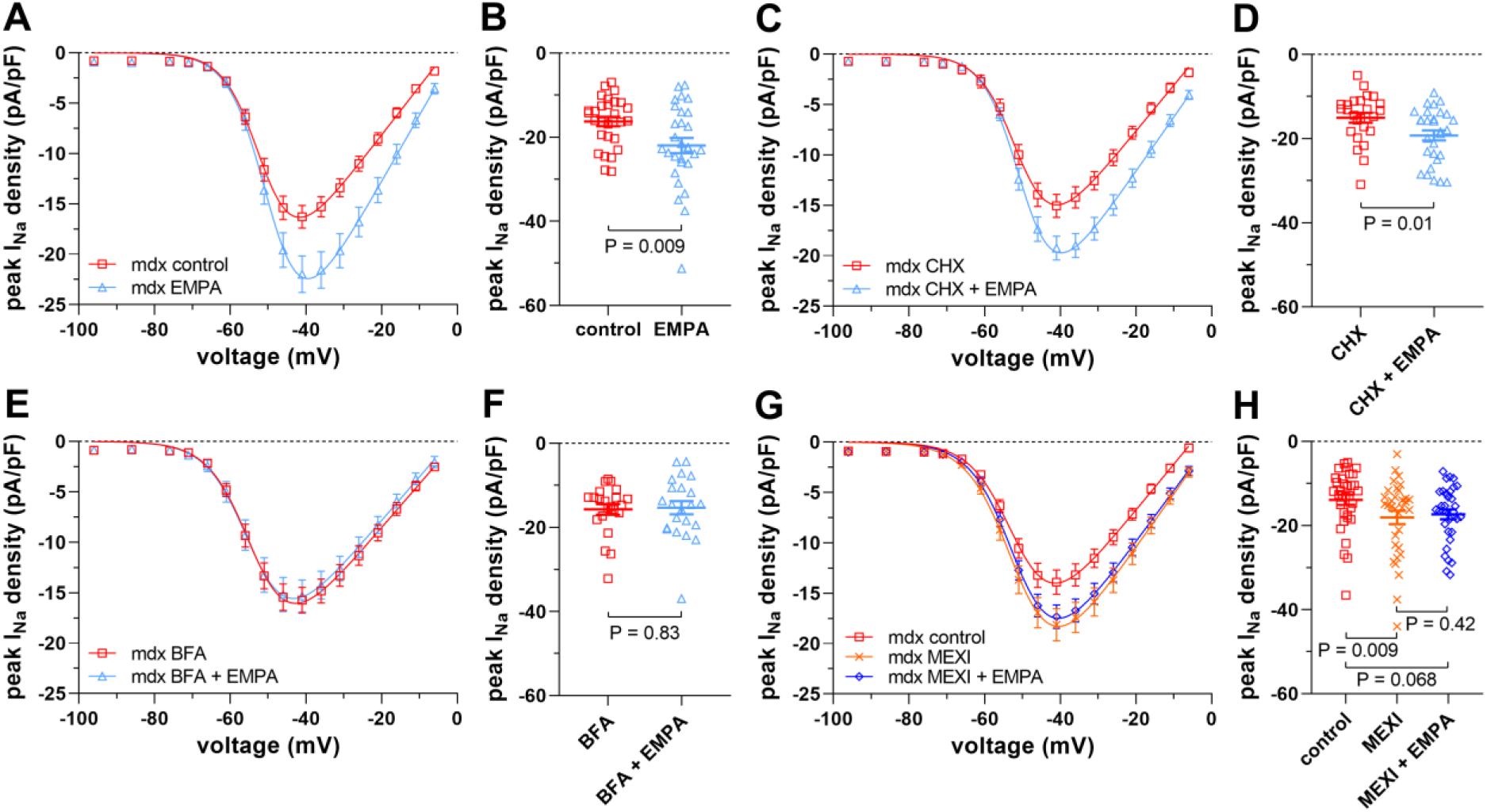
EMPA enhances peak I_Na_ densities in dystrophin-deficient mdx ventricular cardiomyocytes via facilitation of Na_v_1.5 trafficking to the plasma membrane. **A-F**, Effects of protein synthesis inhibition and inhibition of anterograde trafficking on peak I_Na_ enhancement by 1 µM EMPA in mdx cardiomyocytes. **A**, Peak I_Na_ density-voltage relationships from untreated control mdx cardiomyocytes [n = 28 cells], and mdx myocytes incubated for 4 h with 1 µM EMPA [n = 29 cells, all cells originating from the same 4 mdx hearts]. Parameters for I_Na_ activation are given in Table 1. **B**, Dot plot comparing the maximum peak I_Na_ densities of untreated control and 4 h EMPA-treated mdx cardiomyocytes at −41 mV. A significant difference existed between control and EMPA-treated cells. **C**, Peak I_Na_ density-voltage relationships from mdx cardiomyocytes treated with 50 µg/ml of the protein synthesis inhibitor cycloheximide (CHX) for 4 h [n = 25 cells], and mdx myocytes incubated for 4 h with 50 µg/ml CHX and 1 µM EMPA [n = 29 cells, all cells originating from the same 4 mdx hearts]. **D**, Respective dot plot (at −41 mV) showing a significant difference between only CHX- and CHX-plus EMPA-treated cells. **E**, Peak I_Na_ density-voltage relationships from mdx cardiomyocytes treated with 5 µg/ml of the anterograde trafficking inhibitor brefeldin A (BFA) for 24 h [n = 22 cells], and mdx myocytes incubated for 24 h with 5 µg/ml BFA and 1 µM EMPA [n = 22 cells, all cells originating from the same 3 mdx hearts]. **F**, Respective dot plot (at −41 mV) showing that the presence of BFA abolished peak I_Na_ enhancement by EMPA in mdx cardiomyocytes. **G-H**, Effect of 24 h incubation with mexiletine (MEXI) on peak I_Na_ of ventricular cardiomyocytes derived from mdx mice. **G**, Peak I_Na_ density-voltage relationships from untreated control mdx cardiomyocytes [n = 34 cells], mdx myocytes treated with 10 µM MEXI [n = 33 cells], and mdx myocytes treated with 10 µM MEXI and 1 µM EMPA [n = 33 cells, all cells originating from the same 5 mdx hearts]. **H**, Respective dot plot (at −41 mV) showing that MEXI treatment significantly enhanced the peak I_Na_ density in mdx myocytes. The additional presence of EMPA had no effect.

Secondly, we compared EMPA’s effect on peak I_Na_ in dystrophic cardiomyocytes with the effects of the local anesthetic drug MEXI, which is known to act as a pharmacological chaperone of Na_v_1.5 channels.^19–21^ We reasoned if EMPA-as MEXI-is a pharmacochaperone of Na_v_1.5, both drugs should exert similar effects on peak I_Na_ in dystrophic cardiomyocytes. Figure 3G-H shows that 24 h incubation of mdx cardiomyocytes with 10 µM MEXI indeed significantly enhanced peak I_Na_ densities in a similar manner to EMPA (compare with Figure 1A-B). The additional presence of 1 µM EMPA did not further enhance peak I_Na_, suggesting that both drugs share the same mechanism of action.

Thirdly, EMPA binds to Na_v_1.5 in the same region as local anesthetics, whereby the residues F1760 and Y1767 play an essential role.^15^ Here, we tested if disruption of drug binding by mutagenesis (i.e. the introduction of F1760A and Y1767A) affects EMPA’s enhancing effect on peak I_Na_ in Na_v_1.5-expressing tsA201 cells. Figure 4A-C shows that Na_v_1.5-F1760A peak I_Na_ densities were moderately enhanced after 24 h incubation with 10 µM EMPA (estimated free concentration: 1 µM). In contrast to wild-type Na_v_1.5 (inset in B), however, this difference was not statistically significant (Fig. 4C). Strikingly, in Na_v_1.5-Y1767A mutant channels, the EMPA effect was completely lost (Fig. 4D-F). Together, these findings suggested that EMPA binding to Na_v_1.5 via residue Y1767 is essential for enhancement of peak I_Na_. Finally, a respective set of experiments was also performed with MEXI. Figure 4G-J shows that the results were similar in that mutation Y1767A completely abolished MEXI’s enhancing effect on peak I_Na_ (Fig. 4I-J). Na_v_1.5-F1760A peak I_Na_ densities, on the other hand, were significantly enhanced after 24 h incubation with MEXI (Fig. 4G-H). Our mutagenesis studies suggested that both EMPA and MEXI must bind to Na_v_1.5 to enhance peak I_Na_. For the effect of both drugs, the interaction with residue Y1767 in Na_v_1.5 is crucial.

**Figure 4.**
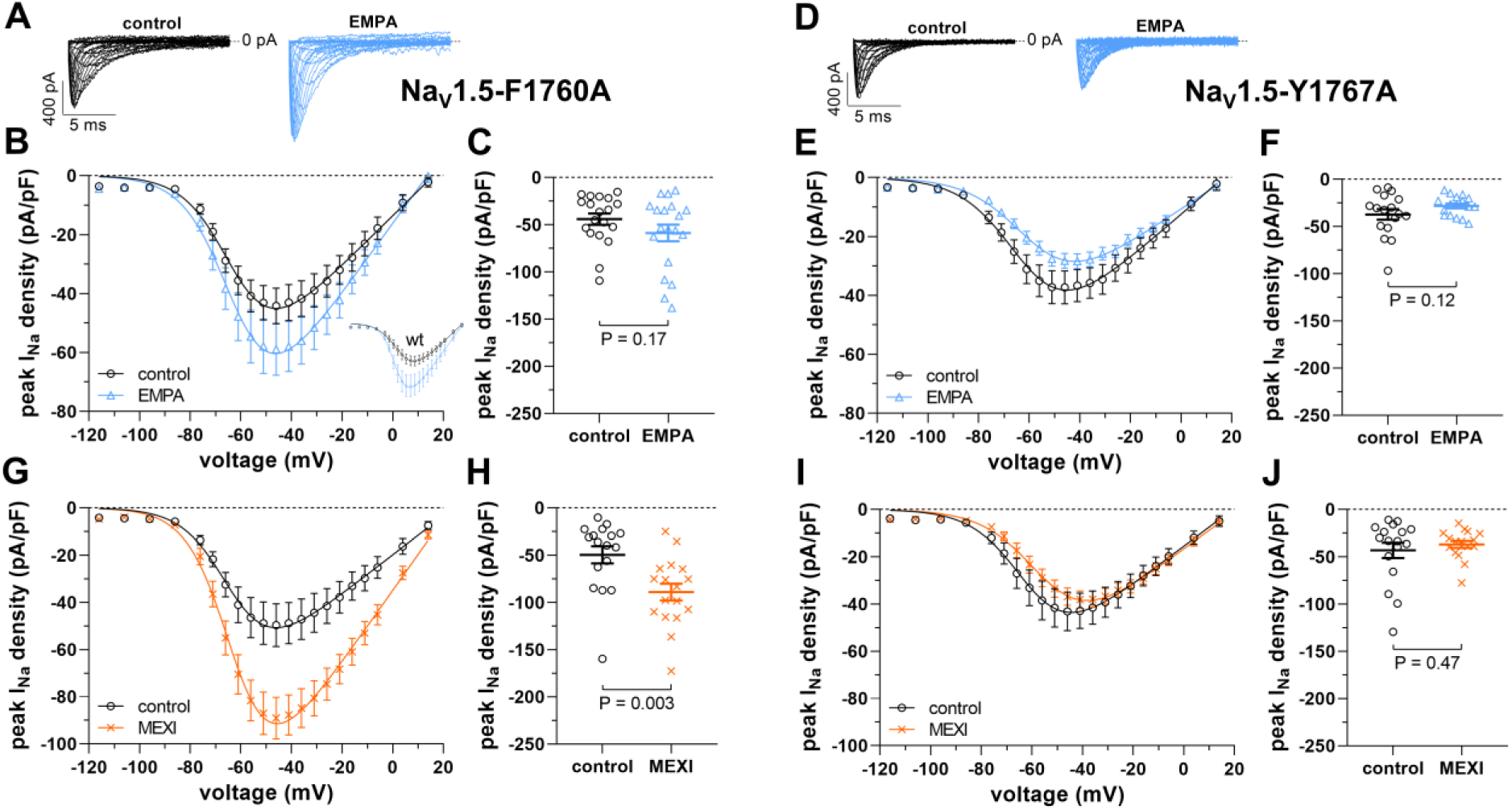
Disruption of EMPA and mexiletine (MEXI) binding to Na_v_1.5 by mutagenesis abolishes the drugs enhancing effect on peak I_Na_. **A-F**, Effect of 24 h incubation with 10 µM EMPA (estimated free EMPA concentration: 1 µM) on peak I_Na_ of tsA201 cells expressing mutant Na_v_1.5 channels. **A**, Typical original current traces of a Na_v_1.5-F1760A-expressing tsA201 cell after incubation under control conditions or after EMPA treatment. **B**, From a series of such experiments [n = 19 cells (control) and 19 cells (EMPA); all cells originating from the same 6 transfections], peak I_Na_ density-voltage relationships were derived. Parameters for I_Na_ activation are given in Table 1. The inset shows respective peak I_Na_ density-voltage relationships derived from tsA201 cells expressing wild-type (wt) Na_v_1.5 (also see Figure 1E). **C**, Respective dot plot at −46 mV. **D**, Typical original current traces of a Na_v_1.5-Y1767A-expressing tsA201 cell after incubation under control conditions or after EMPA treatment. **E**, Respective peak I_Na_ density-voltage relationships [n = 17 cells (control) and 17 cells (EMPA); all cells originating from the same 4 transfections]. **F**, Dot plot at −46 mV. **G-J**, Effect of 24 h incubation with MEXI on peak I_Na_ of tsA201 cells expressing mutant Na_v_1.5 channels. **G**, Peak I_Na_ density-voltage relationships of Na_v_1.5-F1760A-expressing tsA201 cells after incubation under control conditions [n = 17 cells] or after 24 h incubation with 100 µM MEXI [n = 17 cells, all cells originating from the same 4 transfections]. **H**, Respective dot plot at −46 mV. **I**, Peak I_Na_ density-voltage relationships of Na_v_1.5-Y1767A-expressing tsA201 cells after incubation under control conditions [n = 17 cells] or after 24 h incubation with 100 µM MEXI [n = 17 cells, all cells originating from the same 4 transfections]. **J**, Respective dot plot at −46 mV.

### 3.4 Structural analysis of residue Y1767 and molecular docking

To broaden the understanding of our mutagenesis results, we performed in silico simulations and drug docking experiments. Analysis of the nine available cryo-EM structures of rat and human Na_v_1.5 revealed a highly dynamic orientation of the Y1767 side chain (Figure 5A), in contrast to the rigid orientation of F1760. The orientation of Y1767 varies from cavity-facing (e.g., in PDB:6UZ0, shown in orange) to fenestration facing (PDB:8F6P, shown in dark red). Given the medium resolution of the structures (3.2 to 3.5 Å), the local quality of the high-affinity binding region was assessed using the wwPDB Validation 3D report. The density in this region is poor, and the geometry of the Y1767 side chain exhibits rotameric violations and/or clashes, making its orientation uncertain (see supplemental Table 1 for details). Therefore, the Alphafold3 server was used to generate a model of Na_v_1.5 in a non-conductive, closed/inactivated state, free from geometry violations. Modeling this state to investigate the chaperoning effect of drugs is based on the assumption that the structure during trafficking represents a closed and/or inactive state. This is supported by the fact that almost all cryo-EM structures of Na channels were obtained in such conformations, unless mutations were introduced, suggesting these states are energetically more favorable than open conformations. In all Alphafold3-generated models, the Y1767 side chain is predicted to orient towards the lipid-exposed fenestration (colored green, Figure 5A). The Alphafold model with the highest scores was selected for docking. This model exhibits a tightly closed activation gate with 2-fold symmetry in contrast to most available cryo-EM states, such as 6UZ3 (see Figure 5B). Structural changes leading to a narrower activation gate are induced by domain IV movements. Rotation of the lower half of DIV-S6 leads to the reorientation of side-chain L1772 towards the cavity, narrowing the activation gate by ∼ 3 Å (see Figure 5B and supplemental Figure 1). These conformational changes influence the position of the S4-S5 linker and the N-helix of domain IV, but do not noticeably affect the orientation of the IFM motif (supplemental Figure 1). Importantly, the predicted model orients the Y1767 side chain towards the fenestration, leading to favorable and unique drug interactions.

**Figure 5.**
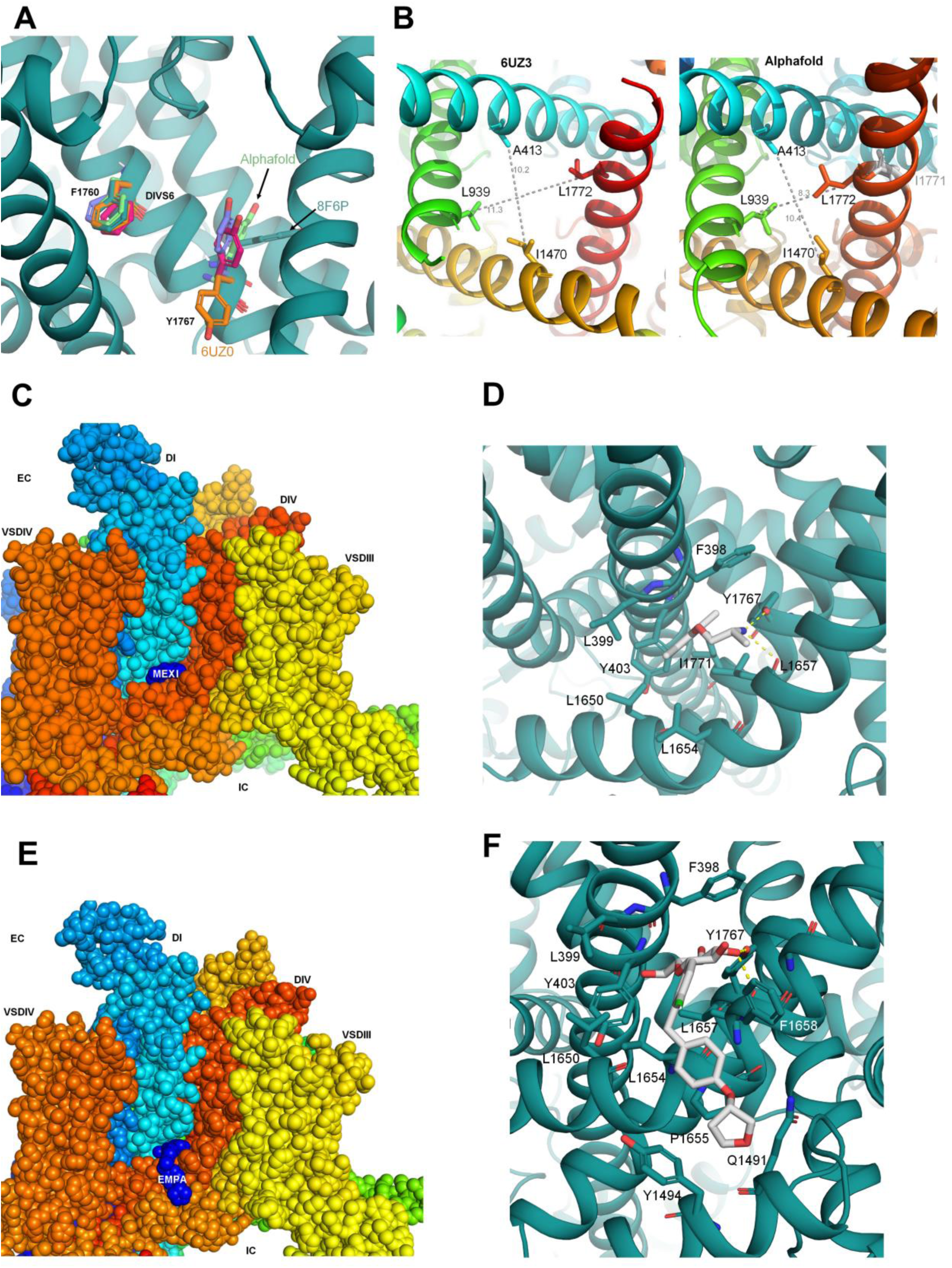
Molecular modeling analysis. **A**, Location of binding residues in different Nav1.5 cryo-EM structures and the Alphafold3 model. The side-chains of F1760 and Y1767 are shown as sticks, with oxygen atoms colored in red. **B**, differences in the activation gate diameter (residue positions taken from Jiang et al.^63^) between opposing C-alpha atoms are shown as gray dotted lines, values in Å. Residue numbering corresponds to human Na_v_1.5 numbers. **C**, spheres representation of the consensus binding mode of MEXI at the fenestration of DI-DIV shown in side view. **D**, close-up view of residues within 5 Å of the drugs, shown as sticks. Hydrogen bonds are shown as yellow dotted lines. **E**, spheres representation of the consensus binding mode of EMPA at the fenestration is shown in side-view. **F**, close-up view of residues within 5 Å of the drugs, shown as sticks. Hydrogen bonds are shown as yellow dotted lines.

Docking of EMPA and MEXI has been previously reported with the goal of identifying the inhibitory site, which is suggested to reside in the fenestrations and the pore of Na_v_1.5.^15,36,37^ In this study, we analyzed the chaperoning effect of these drugs, as well as DAPA and SOTA, to facilitate membrane trafficking. Our functional data revealed an essential role of Y1767 in mediating this effect (Fig. 4); thus, this residue, along with a 10 Å radius, was selected for docking simulations. This selection automatically led to the inclusion of F1760 in the putative binding site, which exerts a moderate functional effect in the case of EMPA but not MEXI (Fig. 4). As shown in Figure 5C-F, MEXI and EMPA exhibit favorable interactions at the DI-DIV interface. All three SGLT2 inhibitors exhibit rather similar binding modes, as shown in supplemental Figure 2 for DAPA and SOTA. The primary stabilizing factor for all drugs is the formation of a hydrogen bond between the hydroxyl group of Y1767 and the primary amine group of MEXI (neutral form) or the sugar moieties of EMPA, DAPA and SOTA. Additionally, EMPA and MEXI are further stabilized by backbone hydrogen bonds to L1657 (DIVS5) and several hydrophobic and aromatic interactions at the interface between DIS6 and DIVS6/S5 in case of SGLT2 inhibitors (see Figure 5E-F, and supplemental Figure 2 for details). No enantiomer-specific differences were observed when docking MEXI. Given the larger size of the three investigated SGLT2 inhibitors, the drugs are predicted to form additional hydrophobic and aromatic contacts mainly with residues L1657 and F1658. While a small number (∼10 %) of drug poses predicted favorable interactions with both Y1767 and F1760 (supplemental Figure 3) in the case of SGLT2 inhibitors, no interactions between MEXI and F1760 were observed, most likely due to the much smaller size of the drug, which precludes simultaneous interaction with both side chains.

## 4. Discussion

Here, we report that 24 h incubation with EMPA significantly increases the peak I_Na_ density of ventricular cardiomyocytes isolated from dystrophin-deficient mdx mice in a concentration-dependent manner with an EC_50_ value of ∼0.1 µM. We thereby confirm respective findings we had previously obtained in the course of both in vivo and in vitro studies.^10^ 1 µM EMPA is considered a therapeutically relevant concentration.^15,16,27^ In the present study, we also demonstrate that EMPA incubation significantly enhances peak I_Na_ densities in dystrophic cardiac Purkinje fibers, tsA201 cells heterologously expressing wild-type Na_v_1.5 channels, and in dystrophic DMD^mdx^ rat ventricular cardiomyocytes. We further show that other SGLT2 inhibitors (i.e. DAPA and SOTA) exert a similar effect on peak I_Na_ in mdx cardiomyocytes as EMPA. By incubating cardiomyocytes for 24 h with EMPA, followed by the removal of the drug prior to recording I_Na_, we made sure to assess the chronic impact of EMPA without a potential acute effect. Apart from that, EMPA at concentrations up to 10 µM neither acutely affected peak I_Na_ in mdx ventricular cardiomyocytes^10^, nor in cardiomyocytes derived from mouse models for heart failure^15,16^ and Na_v_1.5-expressing HEK293 cells.^38^ The Na channel gating properties were also not affected by EMPA.^10,38^ Finally, a chronic peak I_Na_-enhancing effect of EMPA not only occurred in diseased dystrophin-deficient mdx cardiomyocytes (^10^; the present study), but also in mammalian cells heterologously expressing wild-type Na_v_1.5 (^27^; the present study), and in non-diseased human induced pluripotent stem cell-derived cardiomyocytes.^27^ This suggested that EMPA is not only effective in a diseased dystrophic, but also in a non-diseased cellular environment.

### 4.1 EMPA increases peak I_Na_ by pharmacochaperoning of Na_v_1.5

Potential cardiac mechanisms behind the beneficial effects of SGLT2 inhibitors in heart failure, among those modulation of Na_v_1.5 channels, were recently reviewed.^39,40^ Several lines of evidence, listed in the following, strongly suggested that the chronic EMPA-induced peak I_Na_ increase in cardiomyocytes/cell lines can be explained by drug-induced enhancement of Na_v_1.5 channel trafficking to the plasma membrane, i.e. pharmacochaperoning.

First, as demonstrated here by immunofluorescence and confocal microscopy, EMPA-induced peak I_Na_ enhancement in cardiomyocytes goes along with a significantly increased expression of Na_v_1.5 in the plasma membrane.

Secondly, an EMPA incubation time of 4 h-a realistic time frame for protein transport from the ER to the plasma membrane (e.g. see Figure 7 in Málaga-Diéguez et al.^41^) was sufficient to significantly enhance peak I_Na_ in mdx cardiomyocytes (Fig. 3A-B), and BFA, which inhibits the protein transport from the ER to the Golgi apparatus^42^, completely abolished the effect of EMPA. Further, the presence of the protein synthesis inhibitor CHX did not interfere with the EMPA effect. Together, these findings suggested that EMPA releases a certain fraction of Na_v_1.5 channels otherwise trapped in the ER, thereby enabling trafficking to the Golgi apparatus and further to the plasma membrane. The fraction of Na_v_1.5 retained in the ER may indeed be substantial both in heterologous expression systems and in cardiomyocytes.^28^ In accordance with EMPA-induced facilitation of Na_v_1.5 trafficking, EMPA and other SGLT2 inhibitors were previously shown to reduce ER stress, i.e. the accumulation of unfolded proteins in the ER.^43–45^ Enhancement of Na_v_1.5 trafficking by SGLT2 inhibitors may closely resemble facilitation of Na_v_1.8 ER export by the local anesthetic lidocaine in tsA201 cells.^18^

Finally, EMPA-induced peak I_Na_ enhancement in cardiomyocytes/cell lines resembles the respective effects of MEXI, a local anesthetic known to act as a pharmacological chaperone of Na_v_1.5^19–21^, which can diminish ER retention.^46^ In particular, MEXI incubation enhanced peak I_Na_ in mdx cardiomyocytes similar to EMPA, and EMPA did not have an additional effect on peak I_Na_ in the presence of MEXI. Further, both EMPA and MEXI increase peak I_Na_ in mammalian cells heterologously expressing Na_v_1.5 (^19,27^; the present study), and in cardiomyocytes derived from human induced pluripotent stem cells.^20,27^ In addition, both EMPA^15^ and MEXI^47^ bind to the local anesthetic binding site, and its disruption by mutation Y1767A led to a complete loss of both EMPA- and MEXI-induced peak I_Na_ enhancement in Na_v_1.5-expressing tsA201 cells (the present study). These striking similarities prompt us to suggest that both drugs share the same mechanism of action, and that EMPA-just as MEXI-acts as a pharmacological chaperone of Na_v_1.5 channels, thereby increasing their trafficking to the plasma membrane.

Our molecular modeling suggests a state-dependent drug interface between domains DI and DIV to facilitate trafficking of Na_v_1.5. It crucially depends on the orientation of the Y1767 side chain. The orientation of the Y1767 side chain is highly dynamic, as observed in the nine available cryo-EM structures of Na_v_1.5 (Figure 5A). Unfortunately, most structures exhibit steric clashes in this region, complicating the analysis (supplemental Table 1). Nevertheless, it can be concluded that the orientation of Y1767 is influenced by the gating state, with structures exhibiting the narrowest activation gates orienting the Y1767 side chain towards the lipid-exposed fenestrations (e.g., PDB:8F6P, Alphafold model, Figure 5A, B, supplemental Figure 1). When Y1767 is oriented towards the fenestration, it can form transient hydrogen bond interactions that presumably contribute to stabilizing this interface during trafficking (Figure 5C-F, supplemental Figure 2, supplemental Figure 3). Interestingly, this tyrosine residue is unique to the DI-DIV interface. The other three domains have either phenylalanine (in domains II and III) or isoleucine residues at this position, which lack the ability to form hydrogen bonds. This observation aligns with previous molecular dynamics studies^48^, which suggested that the fenestration at the DI-DIV interface is unique. The presence of the tyrosine side chain and other bulky hydrophobic/aromatic side chains at this interface prevents unhindered drug access to the pore via this fenestration when the side chain is more pore-oriented, as seen in most currently available cryo-EM structures.

Molecular modeling accorded well with our functional data in that residue Y1767 was found to be crucial for the effect of both EMPA and MEXI on Na_v_1.5. Further, modeling also predicted very similar effects of EMPA and the other SGLT2 inhibitors tested (Figure 5C-F and supplemental Figure 2, 3). The different effect of mutation F1760A on peak I_Na_ density modulation by EMPA and MEXI (compare Figs 4B and G) can be explained by the complete lack of interaction of MEXI with residue F1760. Compared with MEXI, the affinity of the SGLT2 inhibitors to Na_v_1.5 is expected to be higher due to their ability to interact with both Y1767 and F1760. Generally, their larger size allows for more hydrophobic interactions (compare Figure 5D with 5E, supplemental Figure 2B, D).

### 4.2 Potential therapeutic relevance

Besides other cardiac drugs, current optimal guideline-directed medical therapy for heart failure with and without reduced ejection fraction comprises an SGLT2 inhibitor (EMPA or DAPA).^13^ Among the various beneficial effects of SGLT2 inhibitors on the failing heart discussed^39,40^, their antiarrhythmic properties seem to play a relevant role.^16,49–51^

The effectiveness of SGLT2 inhibitors in DMD cardiomyopathy is unexplored, but a first clinical trial (NCT06643442) has already been initiated. A reduced peak I_Na_, a characteristic feature of cardiomyocytes derived from dystrophin-deficient animal models for DMD^4,5,7–9^, as well as cardiomyocytes derived from induced pluripotent stem cells of DMD patients^2^, slows the action potential upstroke velocity and cardiac conduction, thereby setting the stage for re-entrant arrhythmias and sudden cardiac death.^1^ Consistent with our earlier work^10^, in the present study, we report that chronic treatment with therapeutic concentrations of EMPA rescues peak I_Na_ loss in dystrophin-deficient ventricular cardiomyocytes and Purkinje fibers, the latter cell type being the essential determinant of ventricular conduction velocity. This accords with a shortened QRS interval and an enhanced ventricular conduction velocity caused by EMPA treatment in a mouse model of myocardial infarction.^51^ An EMPA-induced increase in dystrophic Purkinje fiber I_Na_ density by ∼43 %, as observed in the present study (Fig. 1F), is known to speed ventricular conduction.^52–54^ Thereby, the drug may impede arrhythmia development in the dystrophic mdx mouse heart. We speculate that chronic EMPA treatment improves ventricular conduction and diminishes arrhythmia vulnerability in human patients with DMD. EMPA pharmacotherapy also emerges as a potential strategy to enhance peak I_Na_ and cardiac conduction in other arrhythmia disorders associated with reduced peak I_Na_, such as Brugada syndrome^55^. No such pharmacological treatment is currently available.

Based on the findings of the present study, we propose that MEXI may have a similarly beneficial effect on cardiac conduction in patients as EMPA. EMPA is well tolerated by patients^56–59^ and has a very limited propensity to cause drug-drug interactions.^56^ Chronic administration of EMPA may therefore be advantageous compared to chronic MEXI pharmacotherapy. Thus, MEXI inhibits both L-type calcium^60^ and hERG potassium currents^61^, and has gastrointestinal and neurological side effects.^62^

In summary, we provide evidence that EMPA (and probably also other SGLT2 inhibitors) acts as a pharmacological chaperone of Na_v_1.5 channels, and thereby enhances their membrane trafficking. Chronic SGLT2 inhibitor treatment may thereby improve ventricular conduction and diminish arrhythmia vulnerability in human patients affected with DMD and other arrhythmia disorders associated with reduced peak I_Na_.

## Acknowledgments

The authors thank S. Sucic (Medical University of Vienna) and M. Freissmuth (Medical University of Vienna) for very helpful comments and discussions, as well as J. Uhrinova (Medical University of Vienna) for excellent technical assistance.

---

**Supplemental Figure 1.**
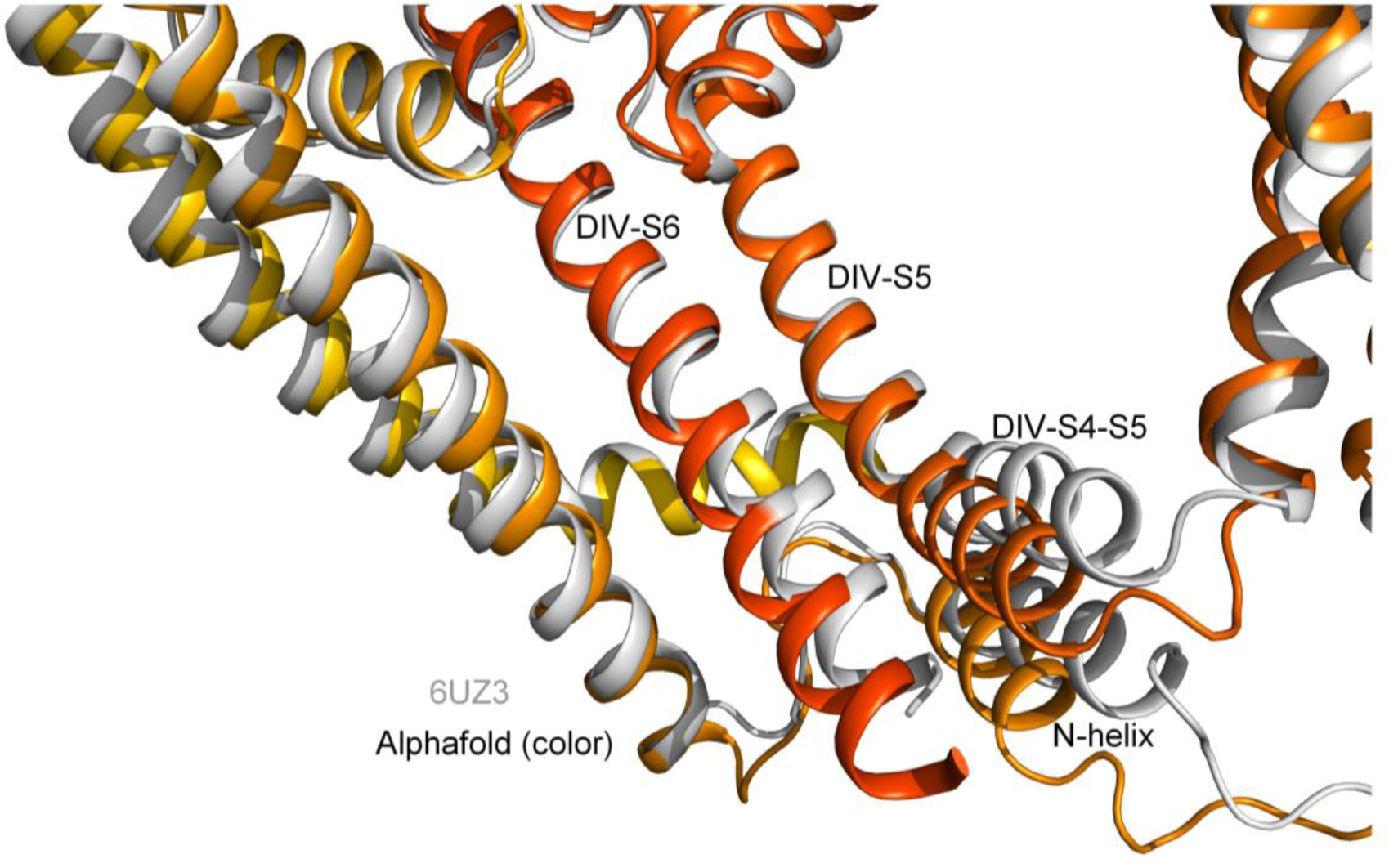
Cartoon representation of DIVS6 and surrounding helices in the Alphafold model shown in color, with the 6UZ3 structure aligned with Swiss-pdb viewer (Guex and Peitsch, 1997; https://pubmed.ncbi.nlm.nih.gov/9504803/) shown in gray. Inward rotation of DIV-S6 is accompanied by movements of the DIV-S4-S5 linker (inward, downward) and the N-helix (inward, downward). Changes in the VSD were not analyzed, since they are not in the vicinity of the drug binding interface.

**Supplemental Figure 2.**
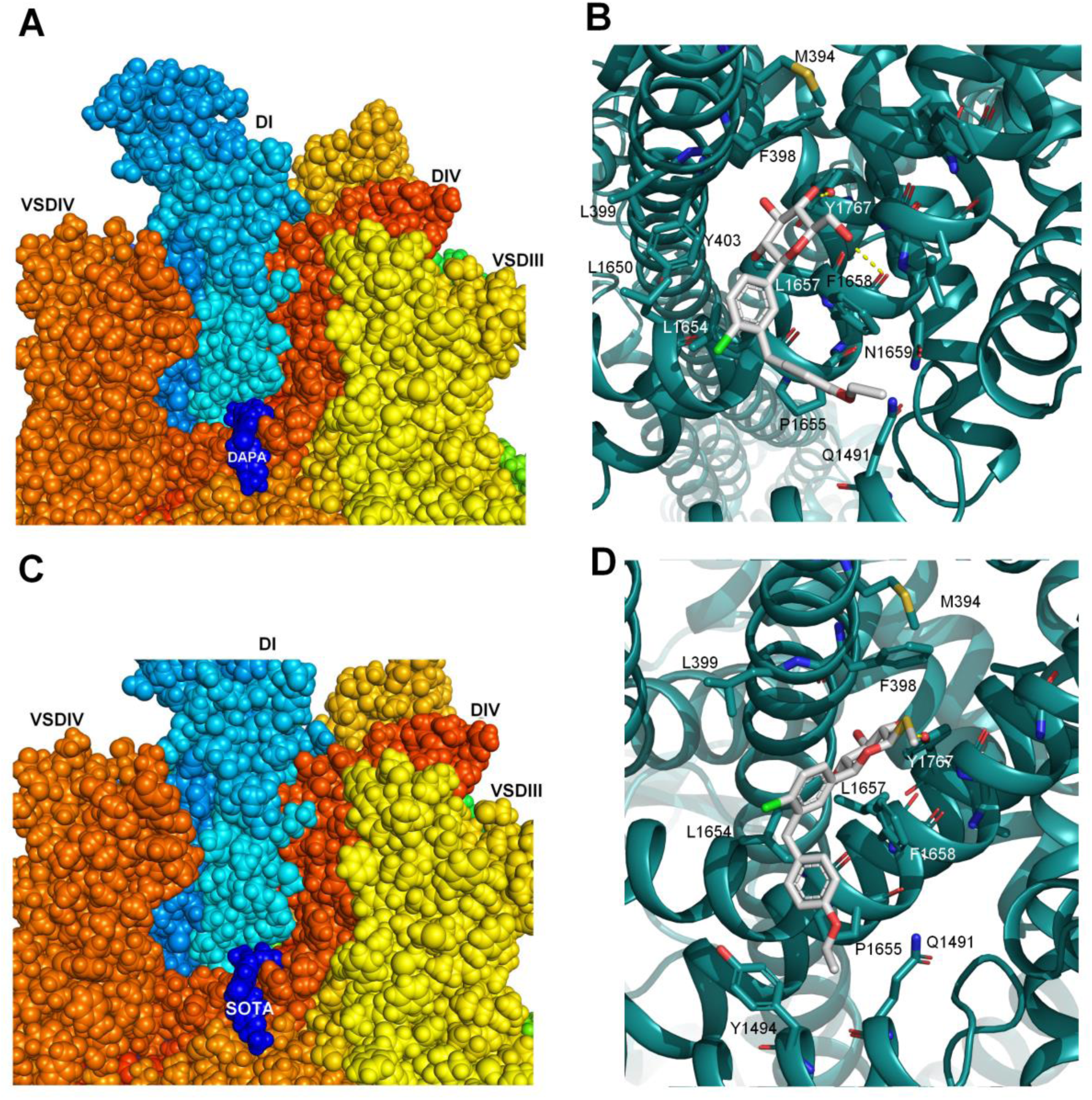
Consensus binding modes of DAPA and SOTA. **A**, **C**, spheres representation of the consensus binding modes of DAPA and SOTA at the fenestration of DI-DIV shown in side view. **B**, **D**, close-up view of residues within 5 Å of the drugs, shown as sticks. Hydrogen bonds are shown as yellow dotted lines.

**Supplemental Figure 3.**
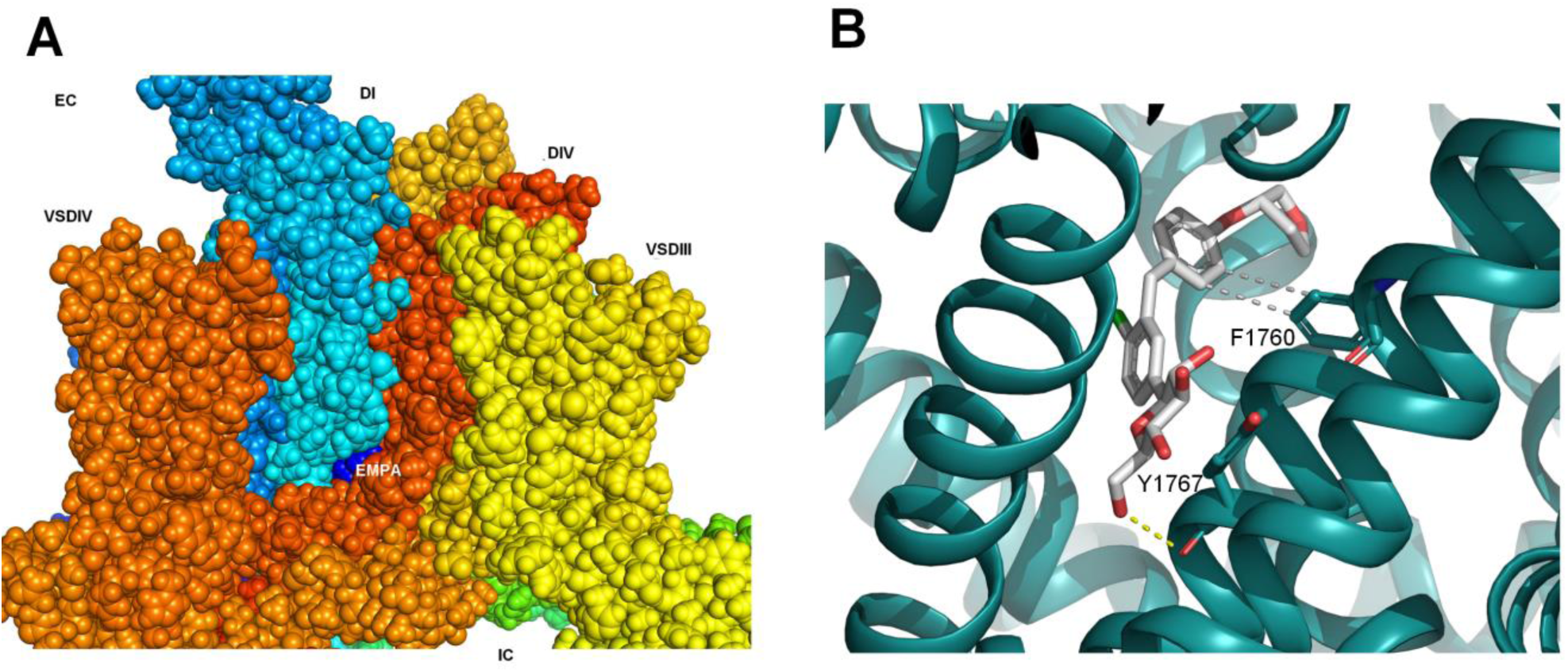
Alternative binding mode of EMPA (∼ 10% of poses). **A**, side view of EMPA binding pose with interactions with F1760 and Y1767 shown in spheres representation. **B**, hydrogen bonds to the backbone of Y1767 side chain are shown as yellow dotted sticks, hydrophobic interactions between the 4-(tetrahydrofuran-3-yloxy) benzyl group of EMPA and the aromatic side chain of F1760 are indicated as gray dotted lines.

**Supplemental Table 1.**
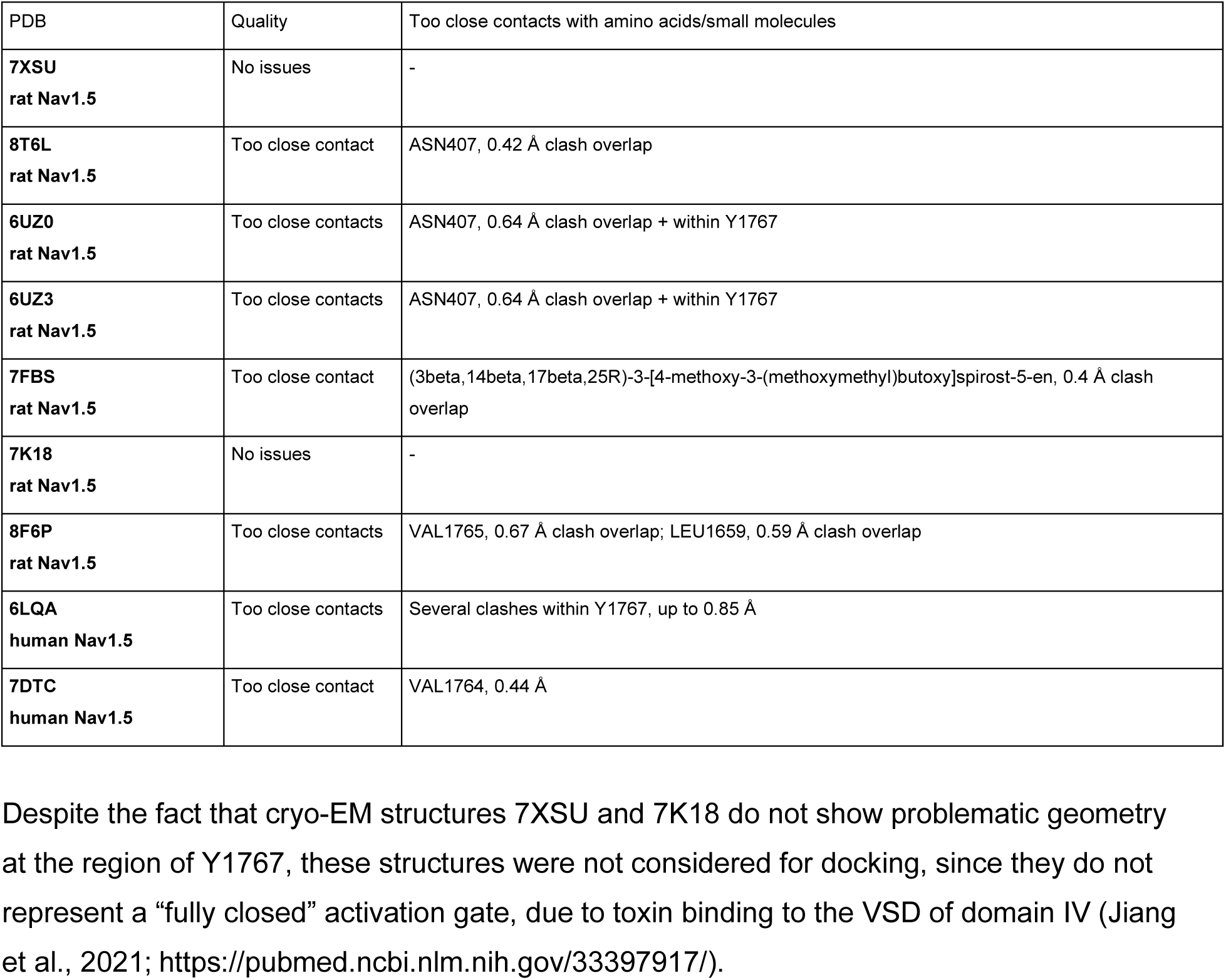

